# GenRank: an R/Bioconductor package for prioritization of candidate genes

**DOI:** 10.1101/048264

**Authors:** Chakravarthi Kanduri, Irma Järvelä

## Abstract

**Summary:** Modern high-throughput studies often yield long lists of genes, of which a fraction are of high relevance to the phenotype of interest. To prioritize the candidate genes of complex genetic traits, our R/Bioconductor package GenRank provides methods that are based on convergent evidence obtained from multiple independent evidence layers. The package facilitates an extensible framework that allows a further addition of novel methods for candidate gene prioritization.

**Availability and Implementation:** The methods are implemented in R and available as a package in Bioconductor repository (http://bioconductor.org/packages/GenRank/).

## 1. Introduction

Genetic studies employ multiple independent lines of investigation spanning panomics approaches to holistically understand the molecular background of complex genetic traits. This includes studying the roles of various forms of genomic variation (e.g. SNPs, InDels, and CNVs) and gene expression (in multiple tissues) and regulation in a single phenotype across single or multiple species (e.g, humans and other relevant model organisms). One of the common characteristic challenges of modern high-throughput experiments across ‐omics fields is that they produce long lists of genes, but only very few of those genes could be of high relevance to the phenotype. Several computational methods have been proposed earlier to prioritize such highly relevant candidate genes (Moreau and Tranchevent, 2012). Besides, metaanalytic approaches that integrate gene-level data from multiple evidence layers have been shown to be successful in identifying and prioritizing candidate genes of complex genetic traits (Ayalew *et al*., 2012). However, no implementation of candidate gene prioritization methods exists in Bioconductor project, which otherwise offers a seamless framework to perform genomic analyses. The majority of the existing meta-analysis related packages on Bioconductor have been exclusively developed to integrate microarray gene expression data, but do not serve the purpose of integrating gene-level data from multiple study-types. Here, we have implemented three methods to integrate gene-level data generated from multiple evidence layers. The evidence layers could be categorized based on several factors like sample-group, study-type, sample-source and so on. Example classifications of evidence layers are shown in Figure 1.

**Figure 1:**
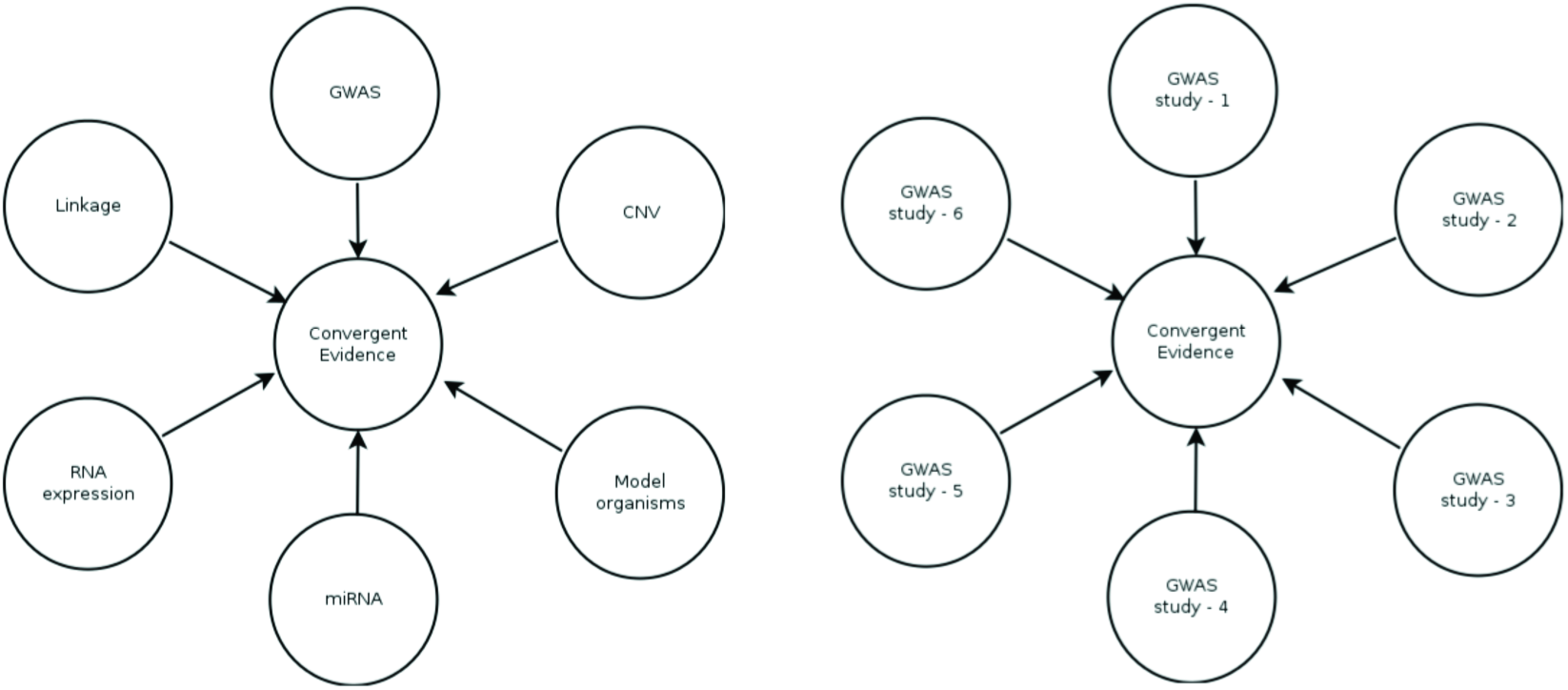
Example classification of evidence layers. Genes can be categorized to different evidence layers based on the type of method used for detection as shown in the left panel. Alternatively, evidence layers could be categorized based on the study material as shown in the right panel. These are just example classification of evidence layers; users can categorize the evidence layers based on several other factors depending upon the research question.

## 2. Implementation

### 2.1 Convergent Evidence (CE) method

Convergent Evidence method is a variant of the famous PageRank algorithm (Page *et al*., 1998). A variant of PageRank algorithm has earlier been adapted to rank genes in microarray-based gene expression experiments (Morrison *et al*., 2005). A conceptually similar gene-level integration has been successfully used to prioritize candidate genes in neuropsychiatric diseases (Ayalew *et al*., 2012).

Here, we modified the PageRank algorithm to compute convergent evidence scores,

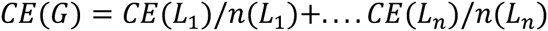

Here *CE*(*L*_*i*_) refers to the self-importance of evidence layer-*i*, while *n*(*L*_*i*_) refers to the number of genes within evidence layer-*i*. Additionally, we propose two other ways to compute convergent evidence scores. One of them is to ignore the number of genes within each layer, thus

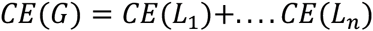

In this case, the convergent evidence score would be equivalent to the primitive vote counting. Another alternative method enables the researchers to determine the importance of each layer based on their own intuition. This involves assigning custom weights to each evidence layer based on their expert knowledge in the field. For example, when a researcher knows that a specific technology could yield less reproducible findings, such evidence layer could be given a relatively less weight compared to the other evidence layers. Another objective way of assigning custom weights to each evidence layer could be based on the sample sizes of each evidence layer. In this case convergent evidence score,

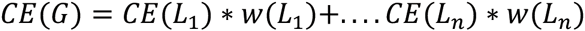

where *w*(*L*_*i*_) refers to the custom weight assigned to evidence layer-i.

### 2.2 Rank Product (RP) method

Rank Product (RP) method has earlier been used widely to perform differential expression and meta-analysis in microarray-based gene expression datasets. This biologically motivated method is quite simple (based on geometric mean) yet powerful to rank genes that are consistently ranked high in replicated experiments (Breitling *et al*., 2004). We adapted the rank product method to identify genes that are consistently highly ranked across evidence layers. In the original rank product method proposed for gene expression datasets, the gene-list or the number of genes (1…*n*) should be the same across replicates/replicated experiments for ranking differentially expressed genes. However, this might not be the case in the majority of instances when integrating gene-level data from multiple evidence layers. For addressing this issue, if there are a total of *n* unique genes across evidence layers, we have added missing genes to each individual evidence layer and gave them a rank of *n* + 1, so that each gene has a rank within each evidence layer. Thereafter, the rank product is computed and compared to a permutation-based distribution of rank product values to estimate the proportion of false predictions (pfp; equivalent to FDR).

### 2.3 Combining p-values

Combining p-values has been one of the traditional methods of meta-analysis. To combine p-values of a gene from multiple evidence layers, the p-values should have been estimated from the same null hypothesis. Popular methods to combine p-values include Fisher's and Stouffer's methods, where the latter incorporates custom weights (e.g. sample sizes). These popular methods have already been implemented in the Bioconductor package *survcomp* (Schröder *et al*., 2011). Here, we built a wrapper around those methods to suit the overarching theme of this package (integrating genelevel data from multiple evidence layers). Missing p-values in some evidence layers could lead to a potential bias when combining p-values. To handle this issue, our implementation returns the combined p-values of only those genes, for which p-values are available at least across 60% of the evidence layers. However, it would be an ideal scenario to have p-values available across all evidence layers.

To avoid a potential bias owing to duplicated genes, duplicated genes are counted only once (as a single vote) within each evidence layer in all the three methods implemented in this package. When retaining duplicated genes, those with significant test statistic (e.g low p-values or high effect-size) were retained.

